# Modeling drug combination effects via latent tensor reconstruction

**DOI:** 10.1101/2021.04.16.439989

**Authors:** Tianduanyi Wang, Sandor Szedmak, Haishan Wang, Tero Aittokallio, Tapio Pahikkala, Anna Cichonska, Juho Rousu

## Abstract

**Motivation:** Combination therapies have emerged as a powerful treatment modality to overcome drug resistance and improve treatment efficacy. However, the number of possible drug combinations increases very rapidly with the number of individual drugs in consideration which makes the comprehensive experimental screening infeasible in practice. Machine learning models offer time- and cost-efficient means to aid this process by prioritising the most effective drug combinations for further pre-clinical and clinical validation. However, the complexity of the underlying interaction patterns across multiple drug doses and in different cellular contexts poses challenges to the predictive modelling of drug combination effects.

**Results:** We introduce *comboLTR*, highly time-efficient method for learning complex, nonlinear target functions for describing the responses of therapeutic agent combinations in various doses and cancer cell-contexts. The method is based on a polynomial regression via powerful latent tensor reconstruction. It uses a combination of recommender system-style features indexing the data tensor of response values in different contexts, and chemical and multi-omics features as inputs. We demonstrate that *comboLTR* outperforms state-of-the-art methods in terms of predictive performance and running time, and produces highly accurate results even in the challenging and practical inference scenario where full dose-response matrices are predicted for completely new drug combinations with no available combination and monotherapy response measurements in any training cell line.

**Availability and implementation:** *comboLTR* code is available at https://github.com/aalto-ics-kepaco/ComboLTR

## 1 Introduction

Combination therapies, involving two or more drugs, offer several advantages over standard monotherapies, including higher treatment efficacies and overcoming resistance mechanisms by modulating multiple targets and signalling pathways. This is especially important in combating complex multifactorial diseases, such as cancer, and cardiovascular, neurological and autoimmune disorders. Moreover, drugs in combination can often be administered in lower individual doses which, in turn, results in reduced risk of adverse reactions (Pemovska *et al*., 2018; Al-Lazikani *et al*., 2012). The number of US Food and Drug Administration (FDA)-approved drug combinations has been continuously growing since the first approvals for co-administration of drugs to treat nervous and respiratory system disorders in 1940s (Das *et al*., 2018). Currently, most of the ongoing research and development is focused on combinatorial therapies for different cancer types (Pemovska *et al*., 2018). The development is, however, very challenging as the number of possible pairwise combinations increases very rapidly with the number of individual drugs, not even mentioning the enormous size of the chemical universe that could be explored (Reymond and Awale, 2012).

Computational approaches offer cost-effective means for large-scale, fast, and systematic pre-screening and prioritisation of potential drug combinations for further experimental validation. Most of the machine learning models introduced to date and benchmarked in crowdsourced DREAM Challenge competitions (Bansal *et al*., 2014; Menden *et al*., 2019) focus directly on the prediction of synergistic drug combinations (Preuer *et al*., 2018; Tonekaboni *et al*., 2018; Li *et al*., 2019; Sidorov *et al*., 2019; Yang *et al*., 2020). Nonetheless, modeling the full dose-response matrices of drug pairs offers more in-depth view of their complex response landscapes, and allows to calculate different synergy metrics as a follow-up step. This is important especially for translational applications, where knowledge of optimal dose combination regions is often critical (Ianevski *et al*., 2020; Tang *et al*., 2015).

Here, we introduce *comboLTR*, a new polynomial regression-based framework for modeling anti-cancer effects of drug combinations in various doses. We compare the performance of *comboLTR* to random forest and recently introduced method by our groups, the *comboFM* method (Julkunen *et al*., 2020), using the NCI-ALMANAC dataset (Holbeck *et al*., 2017) generated by the US National Cancer Institute (NCI). Both *comboLTR* and *comboFM* exploit the fact that dose-response matrices of drug combinations can be represented as a higher-order tensor indexed by drugs, drug concentrations, and cell lines. To predict the response values within the data tensor, a highly non-linear polynomial model is needed in order to capture the multi-way interactions. To learn the parameters of multivariate high-order polynomials, tensor factorization approaches are effective. *comboFM* models the drug combination effects by learning latent factors of the tensor using factorization machines that estimate nonlinear target functions using symmetric polynomials and factorized parametrization (Blondel *et al*., 2016). On the other hand, *comboLTR* is based on Latent Tensor Reconstruction (LTR) method (Szedmak *et al.*, 2020) which can be also considered as an alternative of factorization machines which extends the range of functions that can be learned by removal of the assumption of the symmetry imposed on the polynomials. Moreover, due to only linear dependence on the design parameters separately, a straight, gradient-based algorithm can be applied which can also exploit advanced update rules, e.g. ADAM (Kingma and Ba, 2014). As a consequence, *comboLTR* can process much larger datasets than *comboFM*, in the number of both examples and features, with significantly reduced running time.

In summary, this paper makes the following contributions.

- We introduce *comboLTR*, a new time-efficient framework for modeling drug combination responses in cancer cell lines based on a polynomial regression model where the function learning problem is transformed into a tensor reconstruction problem, with the tensor indexed by drugs, drug concentrations, and cell lines. The algorithm implements mini-batch data processing and allows learning complex, highly nonlinear target functions from large scale data sets with a constant memory complexity and linear running time in all important parameters (degree, rank, sample size, number of variables).
- We demonstrate that *comboLTR* provides highly accurate cell line context-specific results under various prediction scenarios, including more challenging and practical settings where dose-response matrix predictions are made for (i) new drug combinations with no available combination response measurements in any cell line, and (ii) when response measurements of individual drugs are also lacking from the training data. Moreover, we show that drug combination synergy scores can be recovered with high accuracy based on the predicted dose-response matrices.
- *comboLTR* can work with large feature sets, including chemical descriptors and multi-omics cell line features, such as gene expression, copy number variation, CRISPR-Cas9 gene knock-outs, and proteomics data.

## 2 Methods

### 2.1 Notation

In the text, ⨂ denotes the tensor product of vectors, ⟨,⟩ is the inner product, and ∥ ∥ is the norm in a Hilbert space 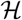. The notation ⟨,⟩ is also applied for the Frobenius inner product of tensors, ○ denotes the pointwise product of tensors with the same shape of any order. 1_*m*_ is a vector of dimension *m* whose all components equal to 1. The set 1, … , *n* for a given *n* is denoted by [*n*]. The matrix **D_u_** is a diagonal matrix whose diagonal is equal to the vector **u**. **A**_*i*_ denotes the row *i* of matrix **A**.

### 2.2 Data representation

In the learning problem, we have a sample of examples given by input-output pairs 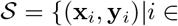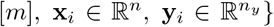 taken from an unknown joint distribution of input and output sources.

The rows of the matrix **X** ∈ ℝ^*m*×*n*^ contain the vectors x_*i*_, and similarly the rows of **Y** hold the output examples, y_*i*_, for all *i*.

### 2.3 Background: learning polynomial regression models

In this paper, we consider learning polynomial regression models

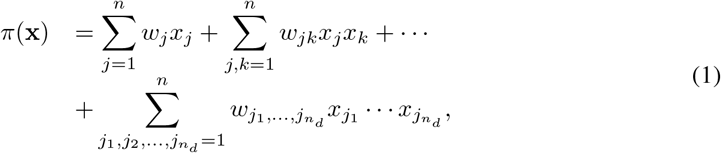

where *w*’s are the regression coefficients to be learned, *n* is the number of input variables and *n*_*d*_ is the degree of the polynomial.

Polynomial regression models are known to have high representation power, capable of accurately representing continuous functions with a fixed *L*_∞_ norm-based t olerance. This fact a llows us to exploit the Stone-Weierstrass theorem and its generalizations, (Prenter, 1970), to approximate those functions by polynomials on a compact subset with an accuracy not worse than a given arbitrary small error.

However, estimating high-degree multivariate polynomial functions presents challenges. An arbitrary multivariate polynomial defined on the field of real numbers can be described by 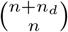 parameters, where *n* is the number of variables, and *n*_*d*_ is the maximum degree of the polynomial. Thus, the complexity relating to the size of the underlying parameter tensor is *O*(*n*^*n*_*d*_^), which grows exponentially in the number of parameters.

This exponential complexity in the polynomial degree presents both statistical and computational challenges: there is often not enough data to reliably estimate all the coefficients, and the exponential time and space complexity forbids processing sufficiently large training sets.

The key approach to tackle this exponential complexity in higher-order factorization machines (HOFM) (Rendle, 2010; Blondel *et al*., 2016) is a special representation of the coefficients as inner products of factors: e.g., for the second order terms

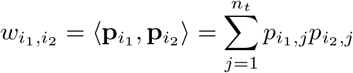

where p_*i*_ ∈ ℝ^*n*_*t*_^ encodes the participation of *i*’th variable in *n*_*t*_ factors. For higher degree terms, the same is given by a generalized inner product

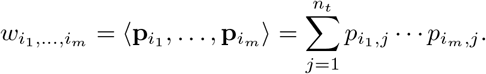

This factorized representation drastically reduces the number of parameters to *O* (*n*_*d*_ · *n*_*t*_ · *n*) (Blondel *et al*., 2016). The HOFM was recently demonstrated to be able to accurately predict drug combination responses (Julkunen *et al*., 2020). However, HOFMs are constrained to symmetric polynomials, that is, functions that are invariant to permutation of features, which restricts the HOFM model as a general regression model.

In this work, we follow an alternative approach for factorizing the parameter representation, called Latent Tensor Reconstruction (LTR) (Szedmak *et al.*, 2020), that starts from the full-order tensor representation of the unknown regression coefficients and learning a factorization into rank-one tensors (of full order) that optimizes the regression error (Table 1). Importantly, the LTR model lifts the limitation of symmetricity of the learned polynomial, and can therefore tackle a wider class of learning problems.

**Table 1:** The general scheme of Latent Tensor Reconstruction-based regression (LTR). Given an *n*_*d*_-order parameter tensor **T** and data point x, the prediction entails computing an inner product between **T** and an *n*_*d*_-order tensor product of the data point with itself. Learning entails finding a factorization of 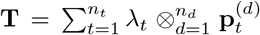 with the lowest regression error.

| Prediction | Learning |
| --- | --- |
| Given : $\mathbf{T}$ tensor, $\mathbf{x}$ | Given : $\{(y_i, \mathbf{x}_i) \vert i \in [m]\}$ |
| Output : $y$ | Output : $\mathbf{T}$ tensor |
| $\pi(\mathbf{x}) = \langle \mathbf{T}, \otimes^{n_d} \mathbf{x} \rangle \Rightarrow y$ | $\min_{\lambda, \mathbf{P}} \sum_i \ y_i - \langle \mathbf{T}, \otimes^{n_d} \mathbf{x}_i \rangle\ ^2$<br>$s.t. \mathbf{T} = \sum_{t=1}^{n_t} \lambda_t \otimes_{d=1}^{n_d} \mathbf{p}_t^{(d)}$ |

### 2.4 Tensor-based representation of polynomial functions

A polynomial function over the real numbers with degree *n*_*d*_ and with *n* variables can also be written in a compact form

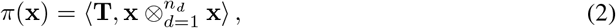

where **T** is a symmetric tensor of order *n*_*d*_, and with dimension 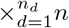 If the vector x is given in homogeneous form, extended with a constant 1, then (2) covers all possible polynomials up to degree *n*_*d*_. The tensor **T** can be given in a decomposed, HOSVD form, (Lathauwer *et al.*, 2000; Kolda and Bader, 2009),

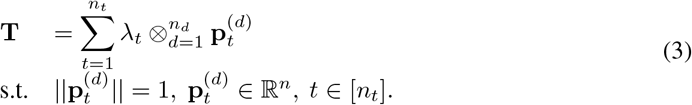

This representation is generally not unique, see (Lathauwer *et al.*, 2000; de Silva and Lim, 2008). By replacing **T** with its decomposed form, the polynomial function turns into the following expressions

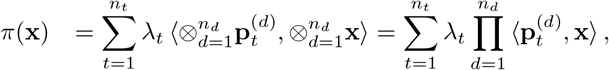

where we exploit the well known identity connecting the inner products and the tensor products (Golub and Loan, 2013). This form only consists of terms of scalar factors, where each scalar is the value of a linear functional acting on the space 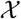. This transformation eliminates the difficulties which arise in working directly with full tensors. Observe that the function *π* is linear in each of the vector-valued parameters, 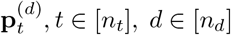.

We can further transform the polynomial representation, (2.4), into a form which does not contain any reference to tensor product. We have a following simple statement, which allows us to introduce an additional factorization within the polynomial function to reduce further the number of parameters.

#### Proposition 1.

The polynomial function *π*(x) can be expressed only by the help of matrix and pointwise, Hadamard, products, namely

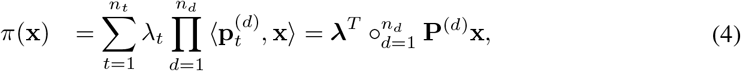

*where* **P**^(*d*)^ *is a matrix of size n_*t*_ × n for any d, whose rows are given as* 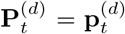, *and* **λ** *is a vector with components λ_t_, t* ∈ [*n*_*t*_].

#### Proof

The matrix-vector product **P**^(*d*)^x yields a vector with components 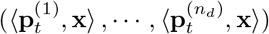, and after a rearrangement, the original form can be restored. ◻

### 2.5 Latent Tensor Reconstruction - basic form

*comboLTR* is built upon the LTR-based polynomial regression method (Szedmak *et al.*, 2020). LTR exploits the representation shown in Proposition 1, which leads to the following optimization problem:

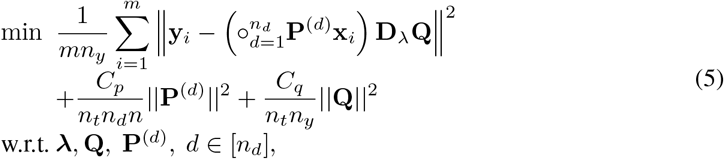

where *C*_*p*_ and *C*_*q*_ are penalty constants, and matrix **Q** projects the vector given by the polynomial function of dimension *n*_*t*_ into the output space.

### 2.6 Reparametrization of the polynomial representation

In the LTR model, the predictor is implemented via a polynomial function *π* acting on vectors x of dimension *n*. The parameter space corresponding to matrices **P**^(*d*)^, *d* ∈ [*n*_*d*_] has dimension *n*_*d*_*n*_*t*_*n* which is large enough to fit the polynomial to a nonlinear function with complex structure, but it requires a large sample to achieve a proper estimation of those parameters. The LTR framework can be extended to increase the flexibility, and in the same time, to reduce the dimension of the parameter space. To this end, let the polynomial function of (4) be reformulated

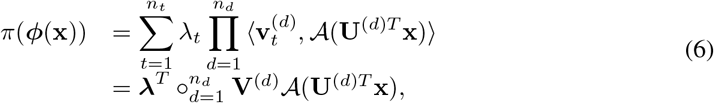

where 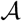 is a pointwise activation function, and the matrix **U**^(*d*)*T*^ is a linear transformation projecting the original input vector into a space with lower dimension, *n*_*k*_, for each *d* ∈ [*n*_*d*_]. That projection can enforce a bottleneck within the polynomial function. This modification preserves the linear dependence on the matrix valued parameters. The expression 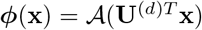 might be viewed as a layer of a neural network. The main difference is that the layers within the LTR are joined by a polynomial function in a parallel way instead of being connected sequentially.

The following table summarizes the matrices describing the extended LTR problem.

|  |  |  |  |
| --- | --- | --- | --- |
| Polynomial | parameters | $\mathbf{V}^{(d)}$ $d \in [n_d]$ | $\in \mathbb{R}^{n_t \times n_k}$, |
| Output | projection | $\mathbf{Q}$ | $\in \mathbb{R}^{n_t \times n_y}$, |
| Scaling | | $\mathbf{D}_\lambda = \text{diag}(\boldsymbol{\lambda})$ | $\in \mathbb{R}^{n_t \times n_t}$, |
| In layers | projection | $\mathbf{U}^{(d)}$ $d \in [n_d]$ | $\in \mathbb{R}^{n \times n_k}$, |
| | scaling | $\mathbf{D}_{\lambda^{(u)}} = \text{diag}(\boldsymbol{\lambda}^{(u)})$ | $\in \mathbb{R}^{n \times n}$, |
| | value | $\mathbf{A}^{(d)} = \mathcal{A}(\mathbf{X} \mathbf{D}_{\lambda^{(u)}} \mathbf{U}^{(d)})$ | $\in \mathbb{R}^{m \times n_k}$, |
| Full | polynomial | $\mathbf{F} = \circ_{d=1}^{n_d} \mathbf{A}^{(d)} \mathbf{V}^{(d)T}$ | $\in \mathbb{R}^{m \times n_t}$, |
| Error | | $\mathbf{E} = \mathbf{Y} - \mathbf{F} \mathbf{D}_\lambda \mathbf{Q}$ | $\in \mathbb{R}^{m \times n_y}$. |

The parameter ***λ*** corresponds to the singular values of the tensor decomposition, see in (3).The extended LTR problem now takes the following form

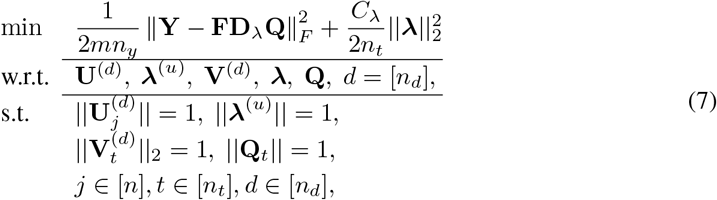

where *C*_*λ*_ is a penalty constant relating to the scale factor ***λ***.

### 2.7 Projection-based algorithm

The optimization problem of (7) is solved by an iterative algorithm which maintains the constrains imposed on the rows of parameter matrices by projecting them onto the unit sphere.

- **Step 1** Let *l* = 0, and the learning speed be 0 < γ < 1.
- **Step 2** Initialize the parameters

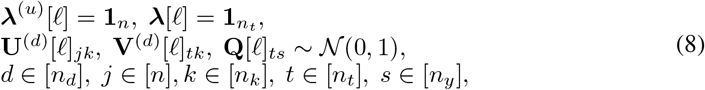

where 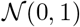 is the standard normal distribution.
- **Step 3** Normalize the rows of the optimization parameters by *L*_2_ norm. Only the vector ***λ***[*l*] will be unnormalized.
- **Step 4** Set the scale value for the unnormalized vector ***λ*** by assuming that all other parameters are fixed. Compute

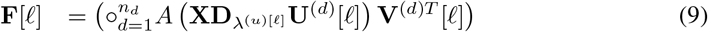

and solve the linear least square problem for ***λ***[*l*]

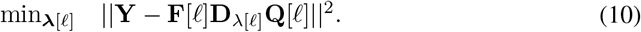
- **Step 5** Compute the value of the objective function of the extended LTR problem given by (7)

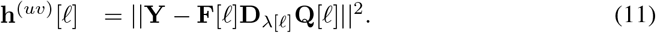
- **Step 6** Compute the partial gradients of h^(*uv*)^ by applying (15),

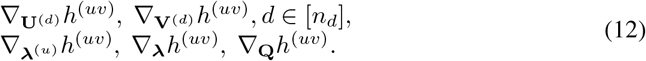
- **Step 7** Update the parameters

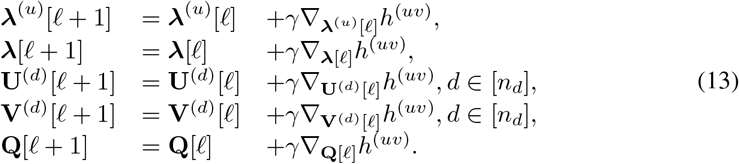
- **Step 8** Normalize the optimization parameters in *L*_2_ norm.
- **Step 9** *l* = *l* + 1.
- **Step 10 Go to** Step 5.

For large-scale applications, the above algorithm is further extended by partitioning the training examples into mini-batches, and processing them sequentially. A single run of the cycle based on mini-batches is taken as an epoch, and repeated. To reduce the variance caused by the partition, a momentum-based update can be applied, e.g., Nestorov Accelerated Gradient method, or the ADAM method frequently applied for Deep Neural Networks (Polyak, 1964; Nesterov, 2005; Kingma and Ba, 2014).

#### 2.7.1 Gradients

Let the matrix **H**^(*d*)^ ∈ ℝ^*m*×*n*_*k*_^ contain the partial derivatives of the activation function with respect to the components of the matrix **XD**_*λ*_^(*U*)^ **U**^(*d*)^, where we exploited that the activation function is a pointwise map of the matrix in its argument. We exploit the following expressions to shorten the gradient formulas

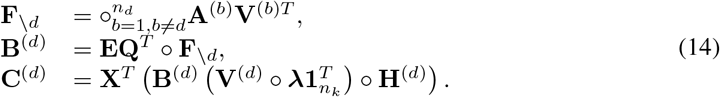

Then, the gradients are expressed in a compact form

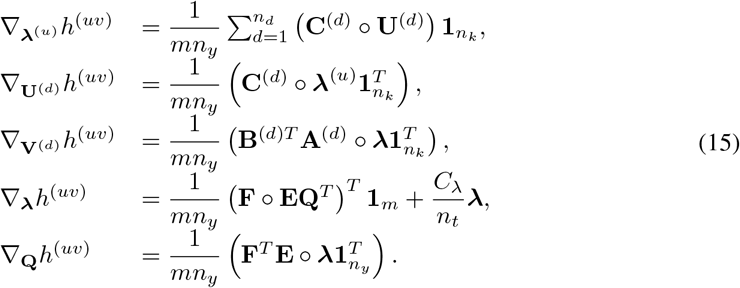

### 2.8 Dataset

In order to evaluate the performance of *comboLTR*, we used the drug combination responses in human cancer cell lines from the NCI-ALMANAC study (Holbeck *et al*., 2017). To exploit more data sources, especially multi-omics profiles of the cancer cell lines, we filtered the data to include only the cell lines for which gene expression, copy number variation (CNV), CRISPR-Cas9 gene knock-outs, and proteomics data were available. The resulting dataset consisted of 828 324 response measurements of 5 035 drug combinations and 15 396 monotherapies in 19 cancer cell lines originating from 9 tissue types. Each drug combination has been screened using 4 × 4 dose–response matrix design. The response measurements are given in the form of percentage growth of the cell lines with respect to a control. The distribution of our drug combination response dataset in 19 cell lines was identical to the distribution of all drug combination responses from the NCI-ALMANAC study, as shown in Figure S1.

### 2.9 Feature representation

Each drug combination response is uniquely determined by five components, that is, two drugs, their concentrations, and a cell line. Such drug combination responses indexed by quintuplets form a fifth-order tensor (Figure 1a). To flatten the higher-order tensor into a feature matrix, each such quintuplet is assigned a unique codeword by one-hot encoding the five components. The resulting tensor index features are similar to ones used in recommender systems (e.g. recommending movies to users). In addition to the tensor index features, the feature matrix also consists of auxiliary features, such as chemical and cell line descriptors, to include more available data sources (Figure 1b).

**Figure 1:**
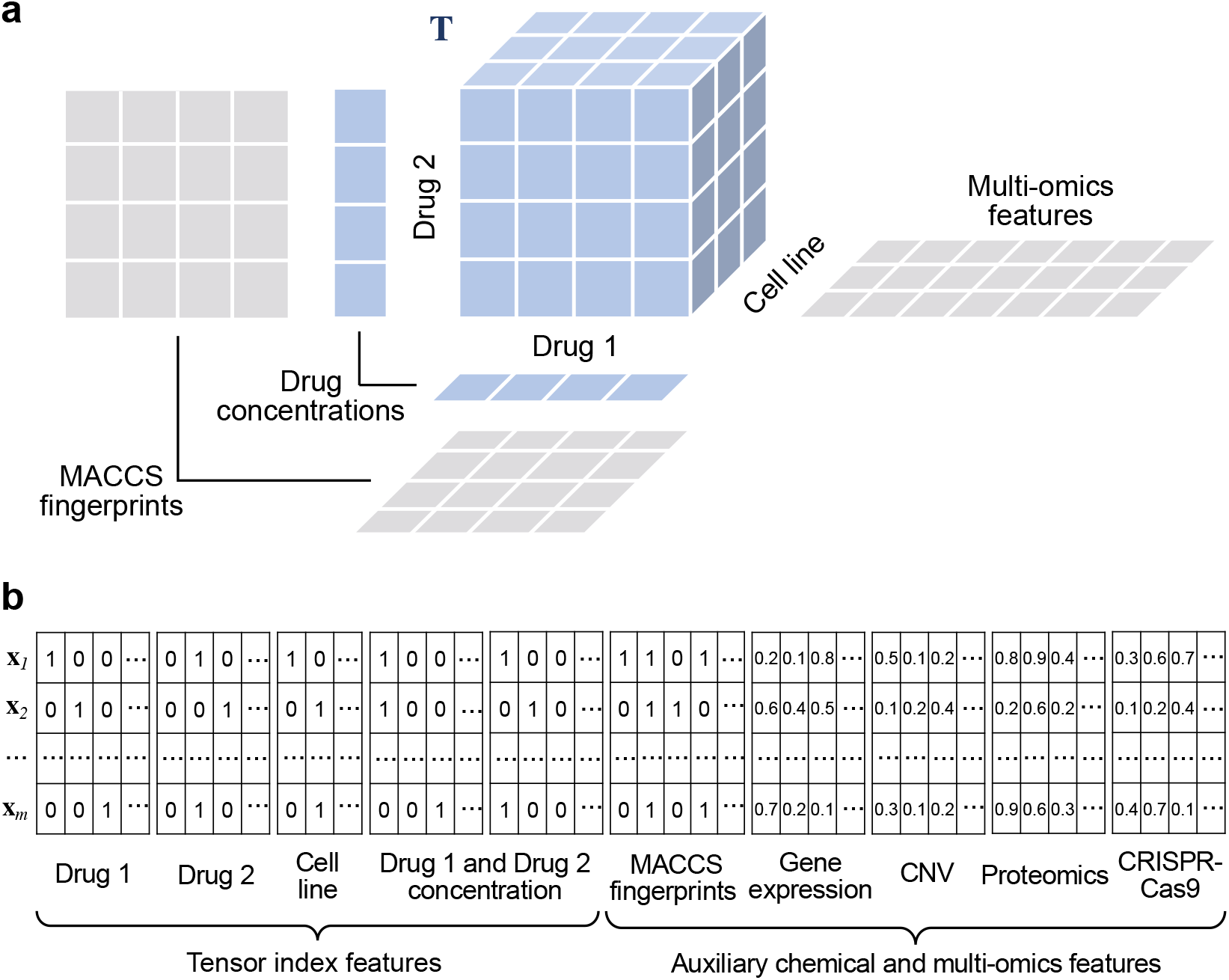
Illustration of the drug combination response tensor and its feature representation. **(a)** drug combination responses form a fifth-order tensor indexed by drugs, their concentrations and the cell lines. **(b)** The drug combination response tensor can be flattened into a tensor index feature matrix via one-hot encoding and accompanied by chemical and biological information.

As for chemical features, we used standard MACCS fingerprint which consists of 166 chemical substructures. Each drug was matched with the substructures and represented as a binary feature vector describing whether a substructure was contained in the drug. Substructures present in all or none of the drugs were removed from the feature set, leaving 148 substructures in the end. For cell line features, multi-omics data including gene expression, copy number variation, CRISPR-Cas9 gene knock-outs, and proteomics data were incorporated from DepMap data portal (Meyers *et al*., 2017; Ghandi *et al*., 2019; Nusinow *et al*., 2020) to represent cell lines. Due to the large size of multi-omics data, in this case more than 70 000 features, only 1% of the omics features with the highest variance across the 19 cell lines were selected, which resulted in 191 gene expression features, 276 CNV features, 174 CRISPR-Cas9 gene knock-out features, and 69 proteomics features.

## 3 Results

We evaluated the performance of *comboLTR* in four practical prediction scenarios (Figure 2):

**Figure 2:**
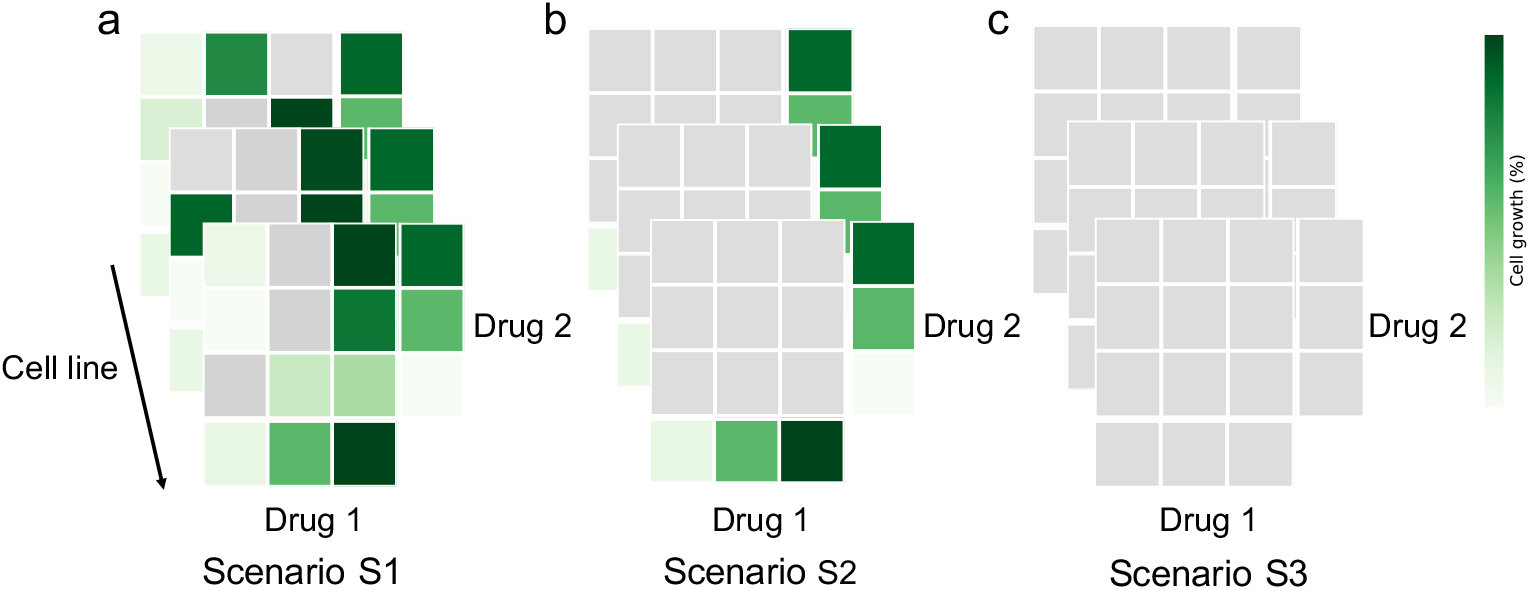
Illustration of different drug combination response prediction scenarios. **(a)** filling in the gaps in partially measured dose-response matrices (S1); predicting dose-response matrices of new drug combinations **(b)** with monotherapy responses (S2) and **(c)** without monotherapy responses (S3).

- Filling in the gaps in partially measured dose–response matrices (S1).
- Prediction of complete dose-response matrices of new drug combinations with no available combination response measurements in any cell line, monotherapy response values present for both drugs (S2).
- Prediction of complete dose-response matrices of new drug combinations with no available combination and monotherapy response measurements in any cell line (S3).

Based on the results by (Julkunen *et al*., 2020), predicting dose-response matrices of completely new drug combinations is the most challenging task from the above. Thus, we aimed to test our *comboLTR* framework under the difficult prediction scenario S2, and furthermore, under even more challenging prediction scenario S3. As shown in Figure 2, S3 has the least information available for a drug combination, since even monotherapy responses of single drugs are not present. Scenario S1 forms a relatively easy task, and thus it was considered as the reference prediction scenario. For completeness, in addition to these three scenarios, following (Julkunen *et al*., 2020), we also considered the scenario of predicting dose-response matrices of previously untested drug-drug-cell line triplets in the case where the response matrices of the same drug combination in other cell lines are known. However, as it was not our main focus in the present study, the results for this prediction scenario are included in the Supplementary material only (Table S1 and Figure S2). We also benchmarked the performance of *comboLTR* against random forest and *comboFM*.

### 3.1 Model optimization via 5-fold cross validation

We applied 5-fold cross validation (CV) in all prediction scenarios in order to tune the model parameters and evaluate the predictive performance. In scenario S1, for each dose-response matrix, combination responses were randomly selected into test sets. In scenario S2, dose-response matrices of certain drug combinations in all cell lines were randomly selected into test sets. All monotherapy responses were kept in the training sets for S1 and S2 prediction scenarios. S3 is similar to S2 but with all monotherapy responses excluded from training set.

Based on our previous research, the degree of the polynomial function in *comboLTR* was set to 5 to model the interactions between the five components, that is, two drugs, their concentrations and the cell line, which uniquely determine each drug combination response in the drug combination response tensor. 20 drugs were randomly selected to subsample the full dataset for *comboLTR* parameter tuning. The subsampled dataset contained 31 095 response measurements of 208 drug combinations and 1 901 monotherapies in 19 cell lines. The subsampled drug combination responses had almost identical distribution as our full dataset and also the whole dataset from NCI-ALMANAC study (Figure S1). Once parameters were determined, the performance of the model was evaluated using 5-fold CV on the full dataset, with the exception that the subsampled dataset was present in the training set only. *comboLTR* was evaluated using 5-fold CV for up to the 9th order with rank 200. Only very slight overfitting was observed in the highest order models.

We used a python implementation of random forest from scikit-learn (Pedregosa *et al*., 2011) and its default parameters for training and prediction. Parameters for *comboFM* were taken from the original publication (Julkunen *et al*., 2020). In addition to fully evaluate the predictive performance of *comboFM* and random forest, in the most challenging prediction scenario S3, their parameters were also optimized using the subsampled drug combination responses. The models with optimal parameters were then applied on the full dataset in the prediction scenario S3.

### 3.2 Prediction of anticancer effects of drug combinations

We used different feature combinations to train *comboLTR* model, as shown in Table 2. Since the one-hot encoded tensor indices are only positional features of the quintuplets, MACCS fingerprint and multi-omics data were used as auxiliary features to provide additional information on drugs and cell lines. In scenario S1, when filling in the gaps in partially measured dose-response matrices, the performance difference between feature combinations was negligible. Using only tensor index features resulted in the Pearson correlation between predicted and measured responses of 0.915, whereas adding auxiliary chemical and biological features led to the Pearson correlation of 0.922. However, in scenarios S2 and S3, when predicting the responses of completely new drug combinations, even without monotherapy responses present, adding auxiliary features clearly increased the prediction performance. In particular, in the most challenging scenario S3, using tensor indices only and additionally including auxiliary features, resulted in the Pearson correlations of 0.893 and 0.915, respectively. With the advantage of handling large feature vectors, *comboLTR* can harness data from different sources for the improved prediction performance. Thus, tensor index features, chemical and multi-omics auxiliary features were used in all further experiments.

**Table 2:** Performance of *comboLTR*, *comboFM*, and random forest (RF) under different prediction scenarios and using different features. Pearson correlations between predicted and measured drug combination responses, reported as averages across 5 cross validation folds ± standard deviations.

| Features | Method | S1 | S2 | S3 |
| --- | --- | --- | --- | --- |
| Tensor indices | *comboLTR* | 0.915±0.009 | 0.894±0.002 | 0.893±0.003 |
| | *comboFM* | 0.920±0.010 | 0.914±0.003 | 0.907±0.004 |
| | RF | 0.886±0.019 | 0.853±0.010 | 0.858±0.010 |
| Tensor indices<br>+ MACCS | *comboLTR* | 0.921±0.010 | 0.908±0.003 | 0.910±0.003 |
| | *comboFM* | 0.923±0.012 | 0.923±0.005 | 0.913±0.005 |
| | RF | 0.921±0.016 | 0.872±0.009 | 0.894±0.005 |
| Tensor indices<br>+ Multi-omics | *comboLTR* | 0.908±0.014 | 0.909±0.007 | 0.911±0.005 |
| | *comboFM* | 0.910±0.027 | 0.904±0.014 | 0.870±0.064 |
| | RF | 0.895±0.019 | 0.859±0.010 | 0.865±0.010 |
| Tensor indices<br>+ MACCS<br>+ Multi-omics | *comboLTR* | 0.922±0.011 | 0.914±0.006 | 0.915±0.005 |
| | *comboFM* | 0.915±0.012 | 0.889±0.024 | 0.878±0.064 |
| | RF | 0.923±0.015 | 0.873±0.009 | 0.896±0.005 |

We used random forest and *comboFM* as comparison methods in all prediction scenarios. Scatter plots of the predicted and measured drug combination responses are shown in Figure 3. As expected, in scenario S1, all three methods achieved comparable prediction performance. Pearson correlations for *comboLTR*, *comboFM* and random forest were 0.922, 0.915 and 0.923, respectively. However, the difference in the performance of the methods became clearly visible in the most challenging and practical scenarios of predicting dose-response matrices of completely new drug combinations with and without monotherapy responses available (S2 and S3). Notably, under scenario S3, *comboLTR*, with a Pearson correlation of 0.915, clearly outperformed *comboFM* and random forest (Pearson correlations of 0.878 and 0.896, respectively). It demonstrates that monotherapies play an important role in predicting higher-order interactions by *comboFM*. On the other hand, *comboLTR* produced more accurate predictions with fewer experimental measurements, which makes *comboLTR* more practical and applicable in recommending combination therapies.

**Figure 3:**
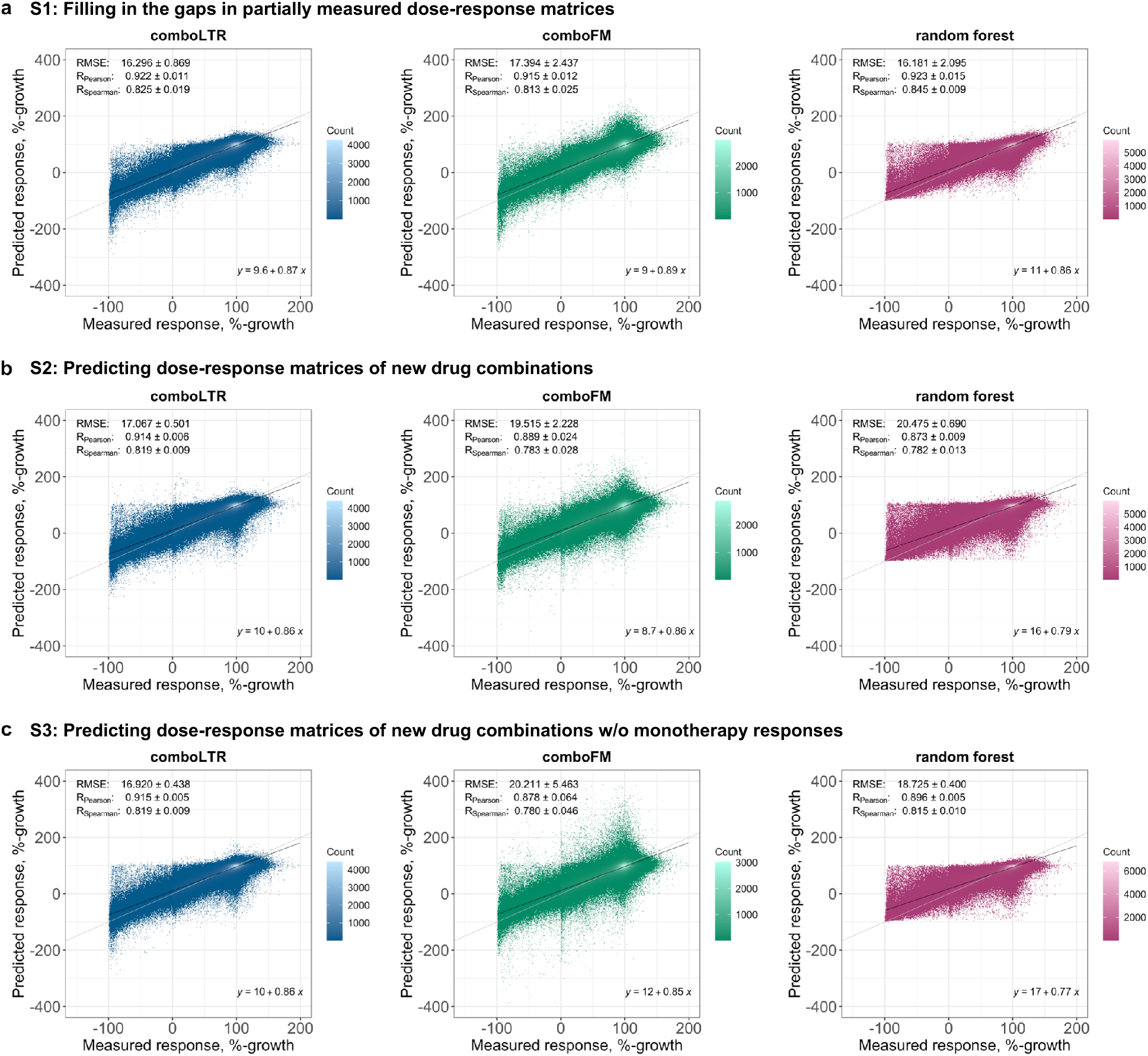
Predictive performance of *comboLTR*, *comboFM* and random forest in three drug combination response prediction scenarios. Scatter plots between the predicted and measured dose-dependent drug combination effects in the form of %-growth of cancer cell lines. The predictions were made under three scenarios of **(a)** filling in the gaps in partially measured dose-response matrices, inferring dose-response matrices of completely new drug combinations with **(b)** and without **(c)** monotherapy responses available. Root mean squared error (RMSE), Pearson correlation and Spearman correlation are reported as averages standard deviations over 5 CV folds. Diagonal line and linear fit are also displayed in each scatter plot. *Note* different x- and y-axes ranges in the plots that are consistent across the panels.

To further study the prediction performance of those three methods, we investigated Pearson correlations for drug pairs in different drug classes and cell lines from different tissue types (Figure 4). In general, *comboLTR* showed higher average Pearson correlation in most tissue types and drug classes. It was also corroborated by the violin plots that *comboLTR* shows better and more stable prediction performance across different drug classes and tissue types, particularly in the more challenging scenarios S2, and even S3 where less information was available.

**Figure 4:**
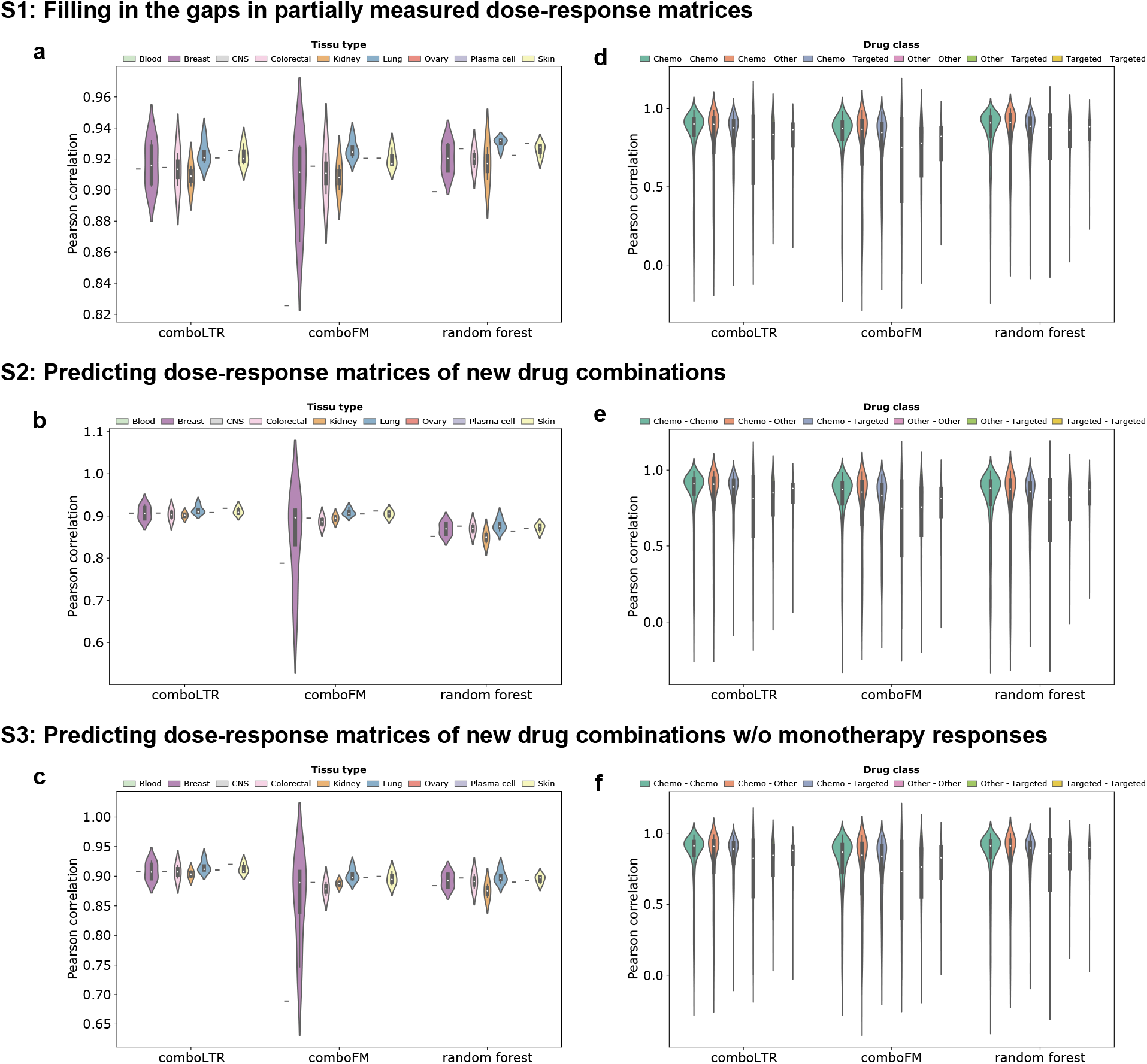
Predictive performance of *comboLTR*, *comboFM* and random forest across tissue types and drug classes in three drug combination response prediction scenarios. Violin plots were used to characterize Pearson correlations of predicted and measured drug combination responses across tissue types **(a-c)** and drug classes **(d-f)**. *Note* that the order of tissue types and drug classes in the legends corresponds to their order in the violin plots.

Next, we evaluated the performance of the methods in quantifying the level of synergy and identifying highly synergistic drug combinations based on the predicted dose-response matrices. To calculate the synergy scores, we applied the NCI ComboScore introduced along with the NCI-ALMANAC dataset (Holbeck *et al*., 2017). Scatter plots and Pearson correlations between the NCI ComboScores calculated based on the complete measured and predicted dose-response matrices of the three methods are shown in Figure S3. Random forest performed particularly well in the simplest scenario S1, but *comboLTR* clearly outperformed the other two methods in the more challenging scenarios, for example, with a Pearson correlation of 0.67 compared to 0.57 (*comboFM*) and 0.46 (random forest), in the S3 scenario. We also conducted discrimination analyses using the precision-recall (PR) curves (Figure S4) and receiver operating characteristic (ROC) curves (Figure S5) to further evaluate the model performance in classifying drug combinations as synergistic vs. non-synergistic with varying thresholds for synergy, in the three prediction scenarios. *comboLTR* showed very competitive performance in discriminating highly synergistic drug combinations across several top-% synergy thresholds in the most challenging prediction scenarios. For example, in scenario S3, the areas under the PR curve (AUPRC) at a synergy threshold of 5% for *comboLTR*, *comboFM* and random forest were 0.25, 0.16, and 0.12, respectively (Figure S4).

To investigate the importance of multi-omics features, contribution of each type of omics data to the model performance was evaluated by “leave one type of omics data out” and “adding only one type of omics data” 5-fold cross validations (Table 3). First, from the feature set comprising tensor indices, MACCS fingerprints and multi-omics data, each type of omics data were excluded to test their contribution to the predictive performance. Then, the predictive performance was also evaluated by including each type of omics data into the feature set, on top of tensor indices and MACCS fingerprints. The prediction performance was relatively stable when including or excluding certain types of omics data.

**Table 3:** The 5-fold cross validation results of *comboLTR*, in the form of Pearson correlations, when leaving one type of omics data out or including only one type of omics data, compared with using the full multi-omics data, on top of tensor indices and MACCS fingerprints.

| Multi-omics feature combination | S1 | S2 | S3 |
| --- | --- | --- | --- |
| Full multi-omics | 0.922 | 0.914 | 0.915 |
| Excluding gene expression | 0.921 | 0.912 | 0.914 |
| Excluding CNV | 0.921 | 0.916 | 0.914 |
| Excluding CRISPR knock-out | 0.921 | 0.914 | 0.913 |
| Excluding proteomics | 0.920 | 0.914 | 0.914 |
| Using only gene expression | 0.921 | 0.912 | 0.911 |
| Using only CNV | 0.921 | 0.911 | 0.910 |
| Using only CRISPR knock-out | 0.921 | 0.909 | 0.911 |
| Using only proteomics | 0.921 | 0.910 | 0.910 |

To investigate the importance of individual features, after the model was trained, each feature column was randomly permuted 20 times, and the average Pearson correlation difference between the models with original and permuted feature matrices was calculated as a measure to evaluate individual feature contribution to the predictive performance (Figure S6). Weights of features from trained *comboLTR* model were also extracted and plotted in the form of a heatmap of L2-norm of each feature set, including MACCS fingerprints, gene expression, CNV, CRISPR-Cas9 gene knock-outs and proteomics data (Figure S6). In general, the most weight has been placed on tensor indices, while MACCS fingerprints and multi-omics data had only a relatively minor contribution to the model accuracy.

## 4 Discussion

Drug combinations are emerging as a powerful treatment modality to combat complex multi-factorial disorders, including cancer. Machine learning models can significantly speed-up the search for effective drug combination therapies by systematically prioritising the most promising combinations for further experimental validation. Using existing drug combination response data, we developed *comboLTR* to efficiently recommend combination therapies for cancer. Table 2 shows that *comboLTR* produces accurate and stable predictions even without monotherapy responses present in the training data. Monotherapy responses, which contain individual drugs tested in different cancer cell lines in various concentrations, are also costly and time-consuming to obtain in the lab. Without such limitation, *comboLTR* is more applicable and practical, especially in clinical research.

We compared the performance of *comboLTR* to random forest and recently introduced *comboFM* method. Both *comboLTR* and *comboFM* are based on polynomial regression. A performance difference between the two methods in prediction scenario S3, where dose-response matrices of completely new drug combinations without monotherapy responses are inferred, could be due to different forms of polynomial functions to be learned. In *comboLTR*, a complete polynomial of degree *n*_*d*_ is used, while in *comboFM*, it is restricted to the symmetric polynomials. Since the monotherapies induce asymmetry in the drug representation, a method which can cover the full range of possible polynomials has higher probability to combine both the symmetric and asymmetric relations. The symmetry restriction of *comboFM* also eliminates the monomials with higher order occurrences of the polynomial variables which could also reduce further the range of functions to be approximated. Thus, it could be hypothesized that *comboFM* may rely more on lower-order interactions, such as monotherapy responses. The lack of such lower-order interaction information results in a performance drop of *comboFM* in the prediction scenario S3. We note that *comboFM* is a bit more competitive without using the MACCS and multi-omics features than when using them, which may be at least in part caused by the symmetricity of the polynomials with respect to the variables. Specifically, the tensor indices and the MACCS and multi-omics features are treated alike by *comboFM*, but not *comboLTR*. Tensor indices, however, do not allow any explanation of the predictions in terms of underlying biological functions or processes, and hence models relying solely on them might not be preferable in practical use.

Furthermore, in practical clinical applications, identification of synergistic drug combinations is of high interest. Importantly, *comboLTR* outperformed *comboFM* and random forest in the most challenging prediction scenarios also in the task of discriminating highly synergistic drug combinations based on the predicted complete dose-response matrices and when using a range of different top-% synergy thresholds (Figures S4 and S5).

Since cross validation experiments were run on different machines, the longest running time and median memory across all prediction scenarios were recorded (Table 4). Compared to *comboFM*, which also learns from higher-order interactions, *comboLTR* is significantly more time-efficient, especially in handling large feature vectors (Table 4). When training models using the full dataset with tensor index features, MACCS fingerprints and multi-omics information, only 3.1 hours were needed for a 5-fold cross validation using *comboLTR*, whereas *comboFM* took 39 hours. Most of the memory was used for storing the data.

**Table 4:** The time (h) and memory (GB) usage of *comboLTR*, *comboFM* and random forest in 5-fold cross validation.

| Features | Time (h) / Memory (GB) |  |  |
| --- | --- | --- | --- |
|  | *comboLTR* | *comboFM* | RF |
| Tensor indices | 1.1 / 24 | 1.2 / 20 | 1.2 / 24 |
| Tensor indices<br>+ MACCS | 2.1 / 35 | 5.9 / 35 | 0.4 / 38 |
| Tensor indices<br>+ Multi-omics | 3.2 / 44 | 31.5 / 63 | 5.6 / 54 |
| Tensor indices<br>+ MACCS + Multi-omics | 3.1 / 74 | 39.0 / 73 | 2.1 / 74 |

In this work, genomic, transcriptomic and proteomic data were used to provide additional information on cancer cell lines. The complex interactions among and within multiple layers of omics measurements form a comprehensive molecular network. For example, CNVs affect expression of genes which, together with post-translational modifications, influence the quantity of proteins. Since the full multi-omics dataset included over 70 000 features, we reduced the dimensionality of the data by selecting only 1% of each type of omics data as auxiliary descriptors characterizing cell lines (see Method 2.7).

As shown together in Table 2, Table 3, and Figure S6, the performance increase brought by integrating multi-omics data into the model was only modest. This phenomenon could be due to that only 1% of the full multi-omics dataset was taken into account, which represented only a small part of the cell’s characteristics. Besides, the complex interactions of the molecular network was not taken into consideration in this experiment. For example, different drugs at various concentrations may result in multiple perturbations to the molecular network, which will change the cellular phenotype into diverse states. The lack of integrating such complicated interactions could lead to less relevant features selected for predicting drug combination responses. One of our future aims is to select cell features based on their connections with tested drug combinations, for example to assign weights to features based on their interaction strength with drugs.

On the other hand, MACCS fingerprint had slightly higher associated feature weights and performance increase, when compared to the multi-omics features. MACCS fingerprint characterizes drugs by the presence or absence of specific chemical substructures. As shown in Figure S7, different drugs in our dataset have common substructures due to the limited number of substructures defined by MACCS fingerprint. Such property is expected to be helpful especially in scenarios S2 and S3 where new drug combinations were predicted. It could be speculated that more weight would be placed on the multi-omics features in the scenario of predicting drug combination responses in the cell lines outside of the training data.

## 5 Conclusions

In this work, we have put forward a novel approach for predicting responses of cancer drug combinations. Our method, *comboLTR*, is based on representing high-degree polynomial regression models through learning a factorization of the parameter tensor containing the unknown regression coefficients. We demonstrated the competitive predictive performance and time efficiency of the *comboLTR* method on the large NCI-ALMANAC dataset (Holbeck *et al*., 2017).

The results indicate that *comboLTR* is a practical tool for prediction and prioritisation of new drug combinations for pre-clinical and clinical evaluation. The ability to predict full dose-response matrices enables a detailed exploration of drug response landscapes and application of different synergy models.

## Acknowledgements

We acknowledge the computational resources provided by the Aalto Science-IT project.

## Funding

This work was supported by Academy of Finland (grants 310507, 313267, 326238, 344698 to T.A.; 313268 and 334790 to J.R.; 311273 and 313266 to T.P.), Cancer Society of Finland (T.A.), Sigrid Jusélius Foundation (T.A.), Helse Sør-øst (grant 2020026 to T.A.), and Doctoral Program in Integrative Life Science (T.W.).

## 1 Supplementary Figures

**Figure S1:**
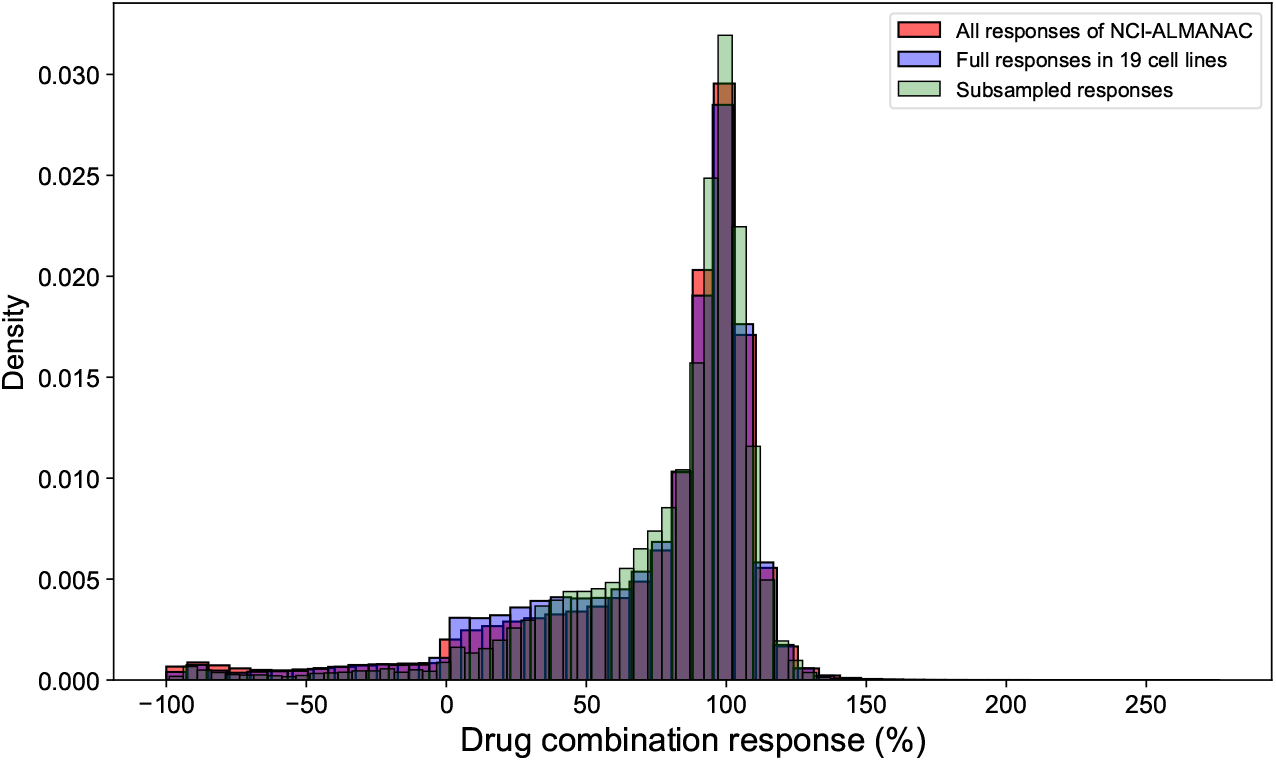
Distributions of drug combination responses of the whole NCI-ALMANAC data; full responses in 19 cell lines; and subsampled dataset.

**Figure S2:**
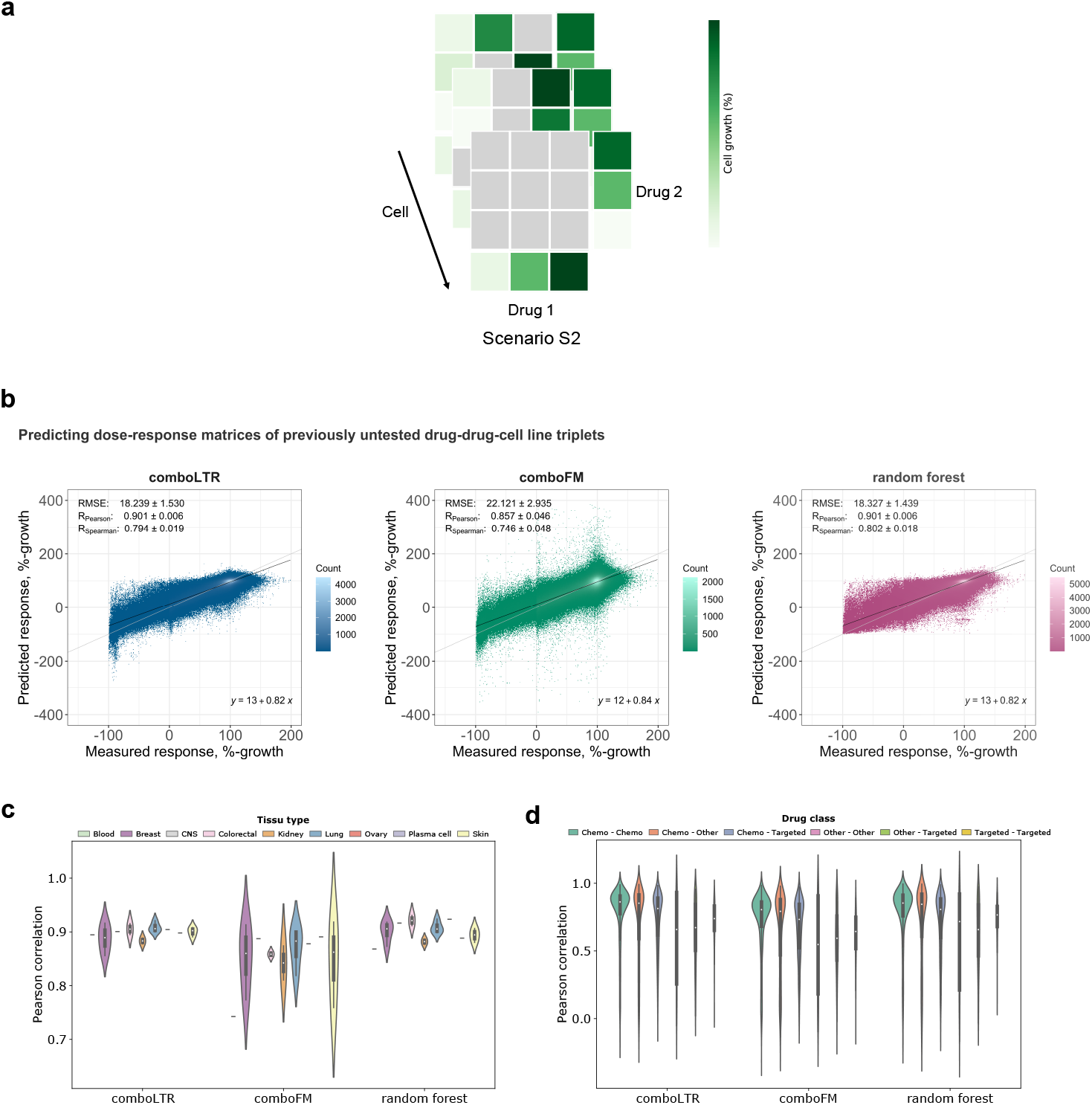
(a) illustration of prediction scenario: predicting dose-responses of previously untested drug-drug-cell line triplets; for each drug combination, the whole dose-response matrices were randomly selected into test sets, such that drug combination is still present in the training set but in other cell lines. (b) predictive performance of *comboLTR*, *comboFM* and random forest in the illustrated scenario: scatter plots of predicted and measured drug combination responses. Prediction performance of *comboLTR* across (c) tissue types and (d) drug classes: violin plots of Pearson correlations of predicted and measured drug combination responses in different tissue types and drug classes.

**Figure S3:**
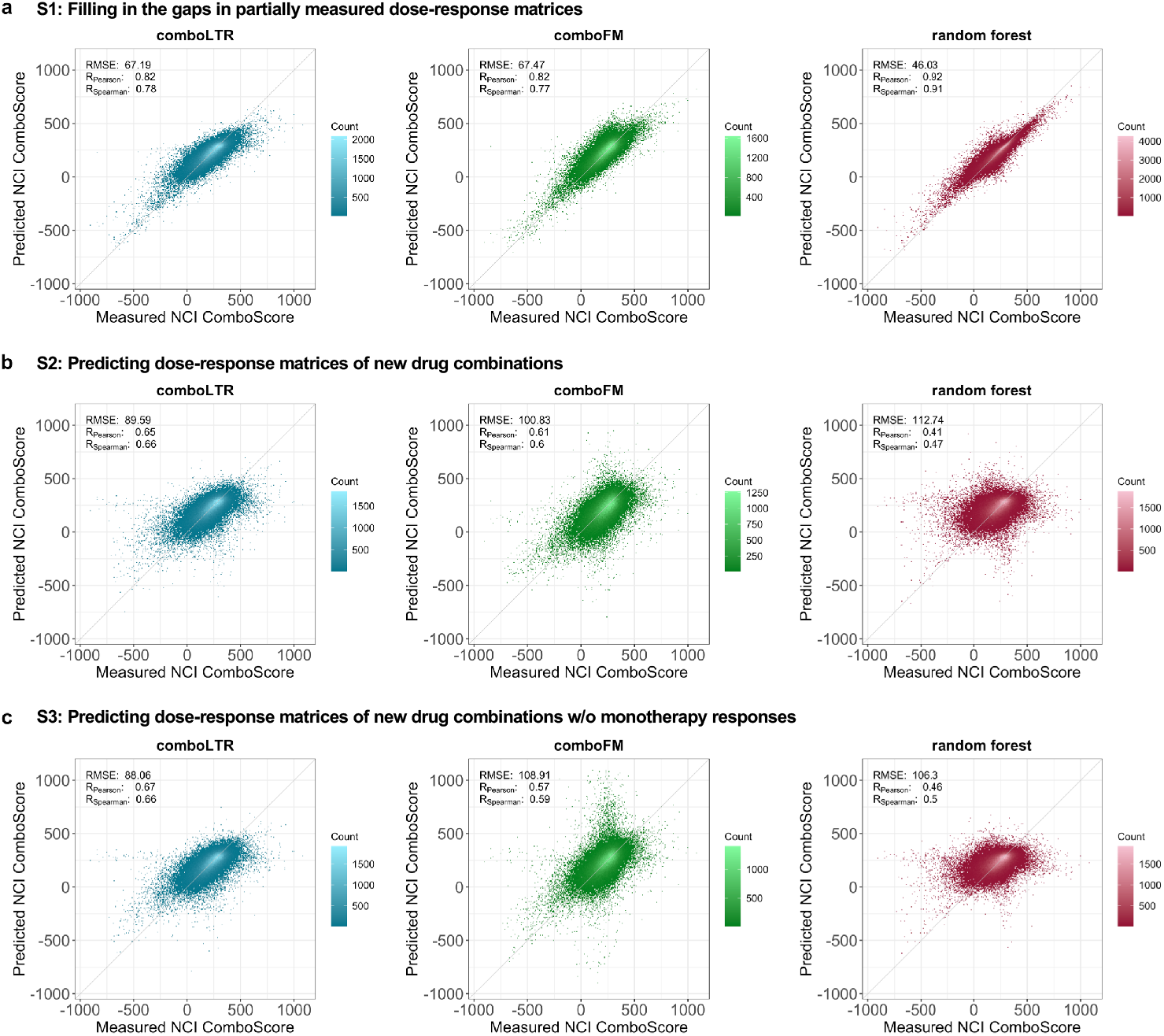
Predictive performance of *comboLTR*, *comboFM* and random forest in predicting NCI ComboScores in three prediction scenarios. Scatter plots between the NCI ComboScores calculated based on predicted and measured drug combination effects in the form of %-growth of cancer cell lines. The predictions were made under three scenarios of (a) filling in the gaps in partially measured dose-response matrices, inferring dose-response matrices of completely new drug combinations with (b) and without (c) monotherapy responses available. Diagonal line is displayed in each scatter plot.

**Figure S4:**
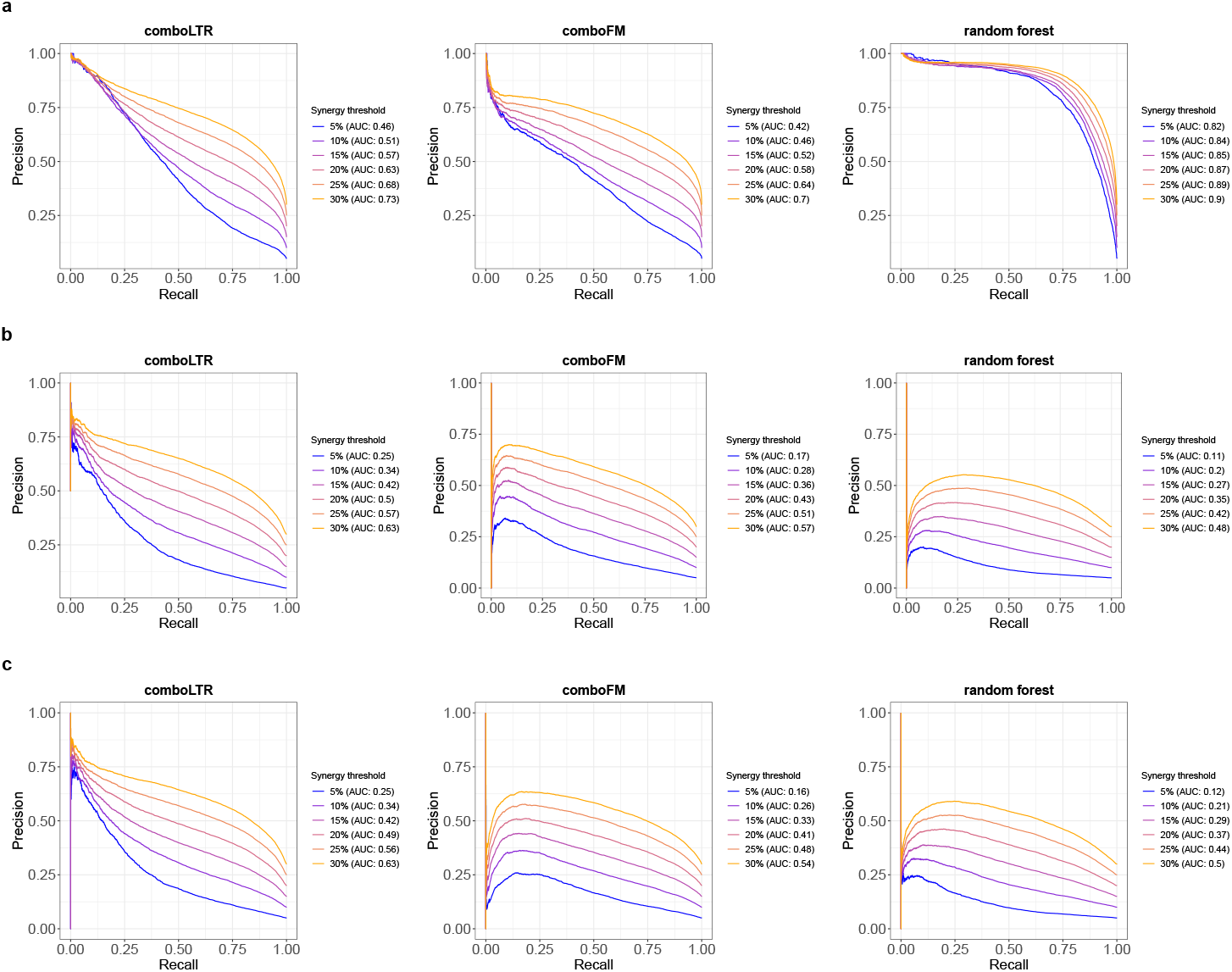
Precision-recall (PR) curves for *comboLTR*, *comboFM* and random forest. PR curves were used to evaluate the model performance in classifying drug combinations as synergistic vs. non-synergistic with varying thresholds for synergy, in the three prediction scenarios: (a) filling in the gaps in partially measured dose-response matrices, inferring dose-response matrices of completely new drug combinations with (b) and without (c) monotherapy responses available. Area under the PR curve (AUPR) is shown in parenthesis.

**Figure S5:**
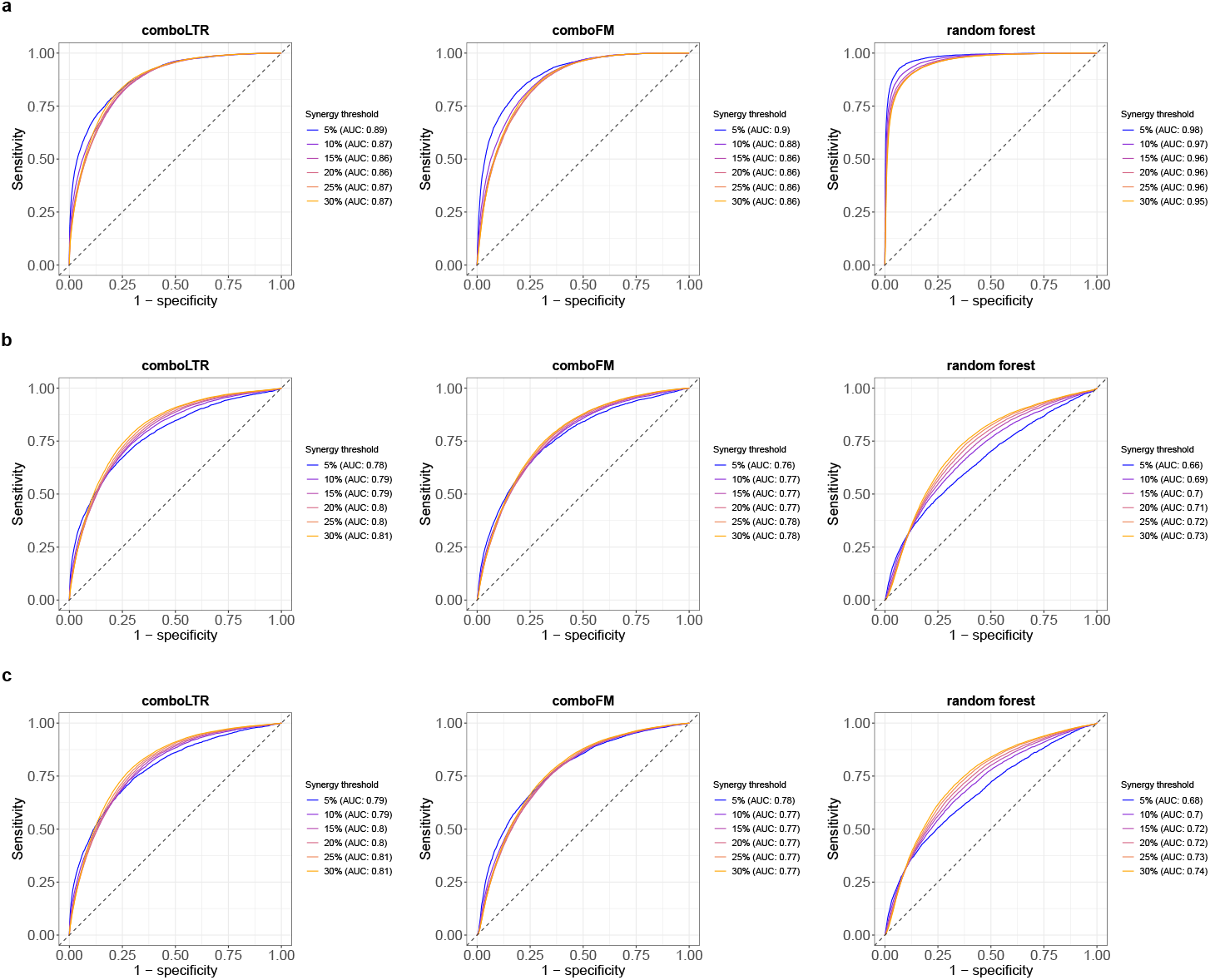
Receiver operating characteristic (ROC) curves for *comboLTR, comboFM* and random forest. ROC curves were used to evaluate the model performance in classifying drug combinations as synergistic vs. non-synergistic with varying thresholds for synergy, in the three prediction scenarios: (a) filling in the gaps in partially measured dose-response matrices, inferring dose-response matrices of completely new drug combinations with (b) and without (c) monotherapy responses available. Area under the ROC curve (AUC) is shown in parenthesis.

**Figure S6:**
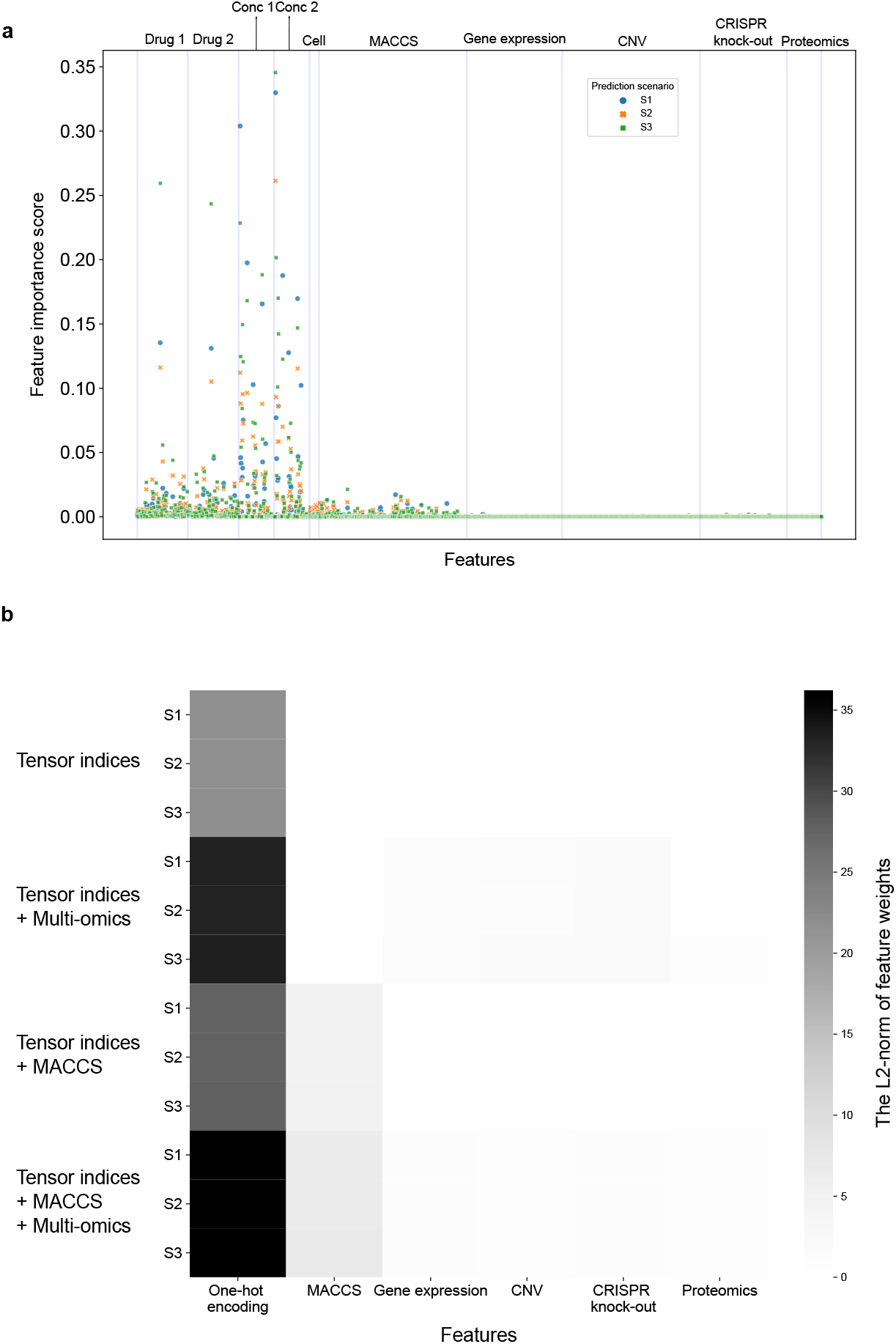
(a) feature permutation importance score of each feature; calculated as the difference of Pearson correlations before and after randomly permuting a feature of all samples. (b) *comboLTR* feature weights in different feature combinations; the L2-norm of the weights of each feature set was used to measure the importance of that feature set.

**Figure S7:**
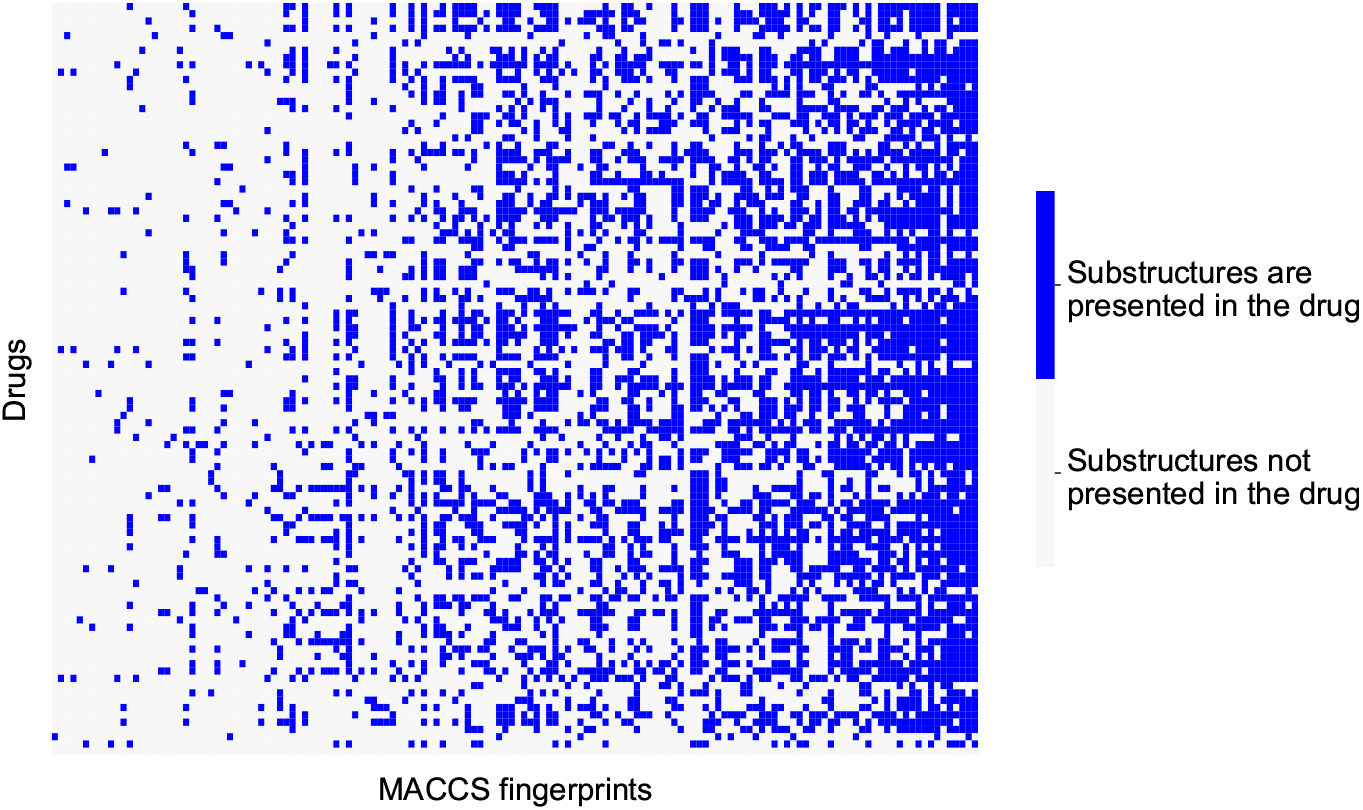
Heatmap of MACCS fingerprints across different drugs. Substructures defined by MACCS fingerprint were matched with drug chemical structures.

## 2 Supplementary Tables

**Table S1:** *comboLTR*, *comboFM*, and random forest Pearson correlations between predicted and measured drug combination responses for prediction scenario: predicting dose-responses of previously untested drug-drug-cell line triplets.

| Feature combination | *comboLTR* | *comboFM* | random forest |
| --- | --- | --- | --- |
| Tensor indices | 0.897±0.007 | 0.904±0.007 | 0.898±0.008 |
| Tensor indices + MACCS | 0.903±0.006 | 0.898±0.045 | 0.899±0.009 |
| Tensor indices + Multi-omics | 0.867±0.012 | 0.884±0.031 | 0.888±0.007 |
| Tensor indices + MACCS + Multi-omics | 0.901±0.006 | 0.857±0.046 | 0.901±0.006 |

## Notes

https://github.com/aalto-ics-kepaco/ComboLTR

### Competing Interest Statement

The authors have declared no competing interest.

